# Post-exposure intranasal IFNα suppresses replication and neuroinvasion of Venezuelan Equine Encephalitis virus within olfactory sensory neurons

**DOI:** 10.1101/2023.06.30.547169

**Authors:** Matthew D. Cain, N. Rubin Klein, Xiaoping Jiang, Robyn S. Klein

## Abstract

Venezuelan Equine Encephalitis virus (VEEV) may enter the central nervous system (CNS) within olfactory sensory neurons (OSN) that originate in the nasal cavity after intranasal exposure. While it is known that VEEV has evolved several mechanisms to inhibit type I interferon (IFN) signaling within infected cells, whether this inhibits virologic control during neuroinvasion along OSN has not been studied. Here, we utilized an established murine model of intranasal infection with VEEV to assess the cellular targets and IFN signaling responses after VEEV exposure. We found that immature OSN, which express higher levels of the VEEV receptor LDLRAD3 than mature OSN, are the first cells infected by VEEV. Despite rapid VEEV neuroinvasion after intranasal exposure, olfactory neuroepithelium (ONE) and olfactory bulb (OB) IFN responses, as assessed by evaluation of expression of interferon signaling genes (ISG), are delayed for up to 48 hours during VEEV neuroinvasion, representing a potential therapeutic window. Indeed, a single intranasal dose of recombinant IFNα triggers early ISG expression in both the nasal cavity and OB. When administered at the time of or early after infection, IFNα treatment delayed onset of sequelae associated with encephalitis and extended survival by several days. VEEV replication after IFN treatment was also transiently suppressed in the ONE, which inhibited subsequent invasion into the CNS. Our results demonstrate a critical and promising first evaluation of intranasal IFNα for the treatment of human encephalitic alphavirus exposures.

**AUTHOR SUMMARY:** Venezuelan Equine Encephalitis virus (VEEV) may enter the brain through the nasal cavity upon intranasal exposure. The nasal cavity normally displays brisk antiviral immune responses, thus it is unclear why this type of exposure leads to fatal VEEV infection. Using an established murine model of intranasal infection with VEEV we identified the initial targets of infection within the nasal cavity and found that antiviral immune responses to virus at this site and during brain infection are delayed for up to 48 hours. Thus, administration of a single intranasal dose of recombinant IFNα at the time of or early after infection improved early antiviral immune responses and suppressed viral replication, which delayed onset of brain infection and extended survival by several days. VEEV replication after IFN treatment was also transiently suppressed in the nasal cavity, which inhibited subsequent invasion into the CNS. Our results demonstrate a critical and promising first evaluation of intranasal IFNα for the treatment of human VEEV exposures.

## INTRODUCTION

Alphaviruses are members of the Togaviridae family of enveloped single-strand RNA arboviruses transmitted by mosquitoes. The arthritogenic Old World alphaviruses include Chikungunya virus (CHIKV), Sindbis virus (SINV), Semliki Forest virus (SFV) and Ross River virus (RRV). The New World alphaviruses, including Venezuelan, Eastern, and Western equine encephalitis viruses (VEEV, EEEV, WEEV), are characterized by the ability to infect the central nervous system (CNS) leading to meningitis and encephalitis, with acute and chronic neurological sequelae [1]. VEEV IAB/IC serotypes are linked to human and equine epizootic outbreaks, while VEEV enzootic cycles occur between rodents and mosquitoes. In addition to natural routes of infection, VEEV, along with EEEV and WEEV, may enter the CNS after intranasal (i.n.) exposure, highlighting the possibility of VEEV weaponization via aerosolization. As there are no approved vaccines for public distribution and no treatments for CNS infection with VEEV, there is a need to understand viral entry and innate immune responses along these routes to develop protective measures.

Studies of murine infections with VEEV enzootic subtype ZPC-738 show that VEEV can enter the CNS through hematogenous spread across an intact blood-brain barrier (BBB) and via anterograde transport along cranial nerves [2,3]. Astrocytes are the first infected cell during hematogenous entry, with further dissemination within the CNS via infected neurons [3]. Intranasal (i.n.) exposure leads to infection of the olfactory sensory neurons (OSN) of the nasal cavity neuroepithelium, leading to neuroinvasion along axons that cross the cribiform plate into the olfactory bulbs (OB), which results in widespread CNS infection and lethality. Low Density Lipoprotein Receptor Class A Domain Containing 3 (LDLRAD3), was identified as a receptor for VEEV [4–6]. Global deletion of LDLRAD3 suppresses systemic infection during the peripheral prodrome phase during which peripheral mononuclear cells become infected [4]. While prophylactic administration of LDLRAD3-Fc fusion proteins suppresses peripheral infection and neuroinvasion, this does not suppress all replication within the brain. It is not known whether LDLRAD3 is expressed by neurons within the olfactory routes of invasion.

Type I interferons (IFN) signal via auto- and paracrine activation of JAK/STAT downstream of the IFN receptor (IFNAR), which is necessary to control initial VEEV infection [7]. However, VEEV has evolved several mechanisms to inhibit IFNAR signaling within infected cells, including host transcription and translation shutoff by VEEV capsid and non-structural protein (nsP)2, and nsP inhibition of IFNAR-induced STAT1 activation via mechanisms independent of host shut off [8–12]. While systemic, pre-exposure (>24 hrs) administration of exogenous IFN controls aerosolized virulent VEEV infection and enhances survival in mice [13], no studies have examined whether IFN has benefit post-exposure. Overall, the multiple routes of entry into the CNS may require specific treatment strategies that depend on the site of initial infection. Alternative to systemic IFN administration, intranasal IFN treatment may uniquely protect the CNS during aerosolized infection. Intranasally administered IFNβ distributes throughout the CNS along olfactory tracts in rats and non-human primates [14,15]. This route of administration resulted in higher concentrations of the cytokine in the brain, suggesting that high doses of IFN may additionally protect susceptible neurons distant from initial sites of neuroinvasion. Intranasal administration of IFNα is well tolerated, making this strategy potentially viable for post-exposure treatment of aerosolized VEEV infection [16].

In this study we demonstrate that VEEV initially targets GAP43+ immature (i)OSN within the olfactory neuroepithelium (ONE). Tropism toward iOSNs correlated with higher LDLRAD3 expression within iOSN versus mature (m)OSN, but no broad deficits in innate immunity, as assessed via scRNAseq, were observed in iOSN that would contribute to their enhanced infectivity over mOSN. Despite rapid VEEV neuroinvasion, host nasal cavity and CNS IFN responses are delayed for up to 48 hours during VEEV neuroinvasion, representing a potential therapeutic window. Thus, we evaluated the efficacy of single dose recombinant IFNα administered intranasally at the time of or early after infection (0-3 hours post-infection), which was able to trigger ISG expression in both the nasal cavity and OB. IFNα treatment delayed onset of sequelae associated with encephalitis and extended survival by several days. VEEV replication after IFN treatment was also transiently suppressed in the ONE, which inhibited subsequent invasion into the CNS. Together these data identify iOSN that express high levels of LDLRAD3 as the initial target of VEEV, define OSN ISG transcriptomic signatures, and demonstrate the efficacy of intranasal delivery of IFNα to protect sites critical to early VEEV CNS infection. Our results demonstrate a critical and promising first evaluation of such a treatment strategy for human encephalitic alphavirus infection.

## RESULTS

### Immature olfactory sensory neurons are the initial site of VEEV infection after intranasal exposure

Intranasal (i.n.) infection with an enzootic strain of VEEV (ID, ZPC-738; herein VEEV) results in rapid progression of weight loss and onset of encephalitic symptoms, with a mean survival time (MST) of 6.5 dpi (S1A Fig). In previous studies we demonstrated i.n. VEEV infection rapidly disseminates into the OB and CTX within 24 hours, with viral loads peaking at 2-3 dpi in the OB and at 4-6 dpi in CTX, hindbrain regions, and spinal cord, highlighting early infection to the OB as a critical period to control VEEV dissemination along the olfactory route after i.n. exposure [3]. To define cellular tropism within the ONE we utilized a reporter strain of VEEV, ZPC-738-GFP, which exhibits similar virulence as the parent strain and labels infected cells green, in conjunction with OSN markers, all detected via double-label confocal microscopy [17]. Within the infected ONE, GAP43^+^ iOSNs are the earliest site of infection at 1 dpi (Figure 1A, B (top) white arrowhead) OMP^+^ mOSN were also infected at this time-point (Figure 1A, (open arrowheads). GFP is also detected within GAP43+ and OMP+ OSN axons traversing the cribriform plate and within the olfactory nerve layer (ONL) of the OB (Fig 1B, bottom). VEEV RNA, as assessed via fluorescent *in situ* hybridization, is also detected within GAP43+ and OMP+ axons within the ONE and OB ONL (Fig 1C). By 3 dpi, GFP+ cells are observed throughout the ONE and OB, including the glomerular, mitral cell, and ganglion cell layers (Fig 1D,E). At this timepoint, VEEV infection continues to spread into the forebrain along the olfactory tract and piriform CTX (Figure 1E). Together, these data indicate that VEEV may utilize both immature and mature OSNs for anterograde transport into the OB.

**Fig 1.**
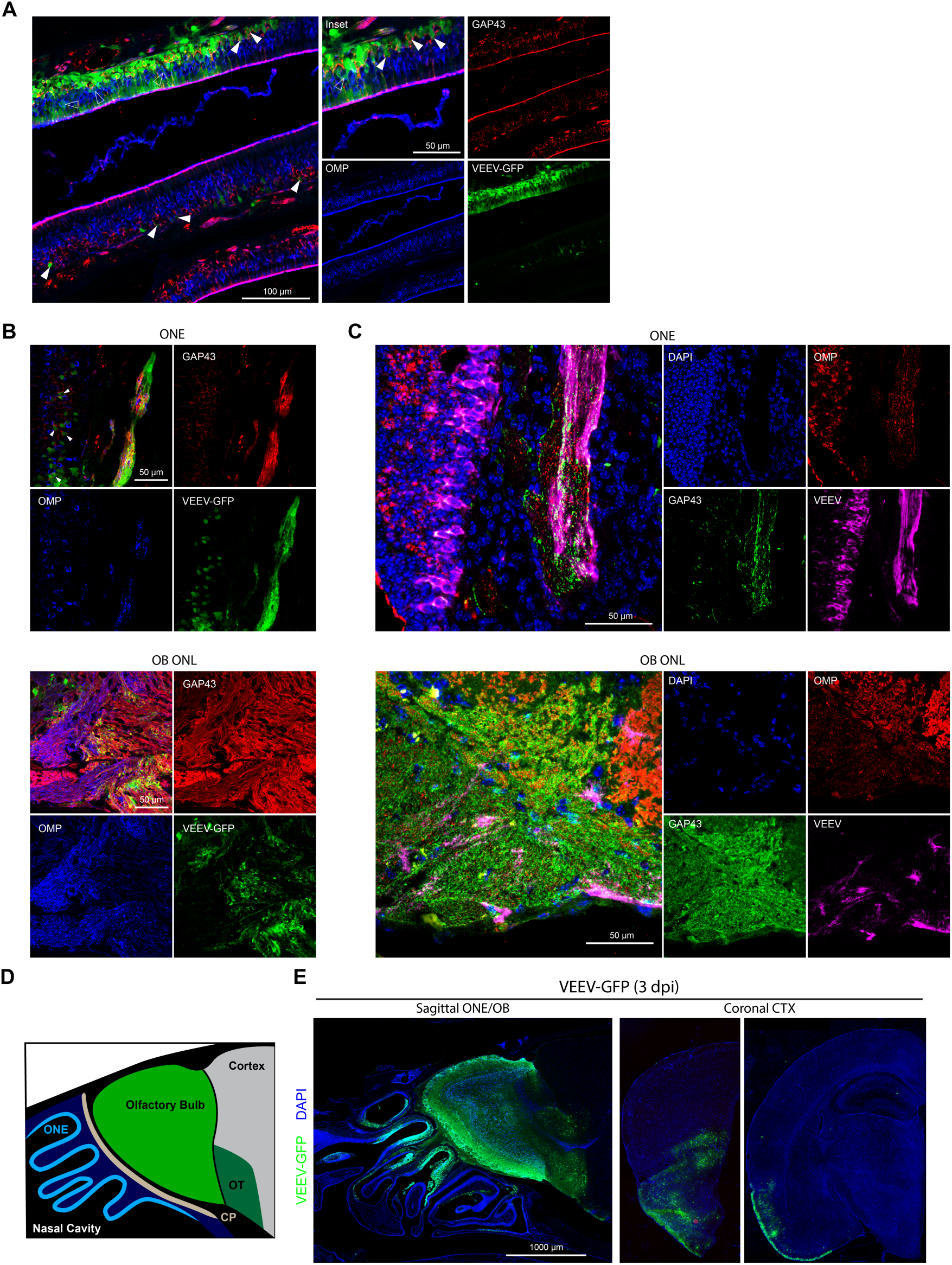
Neuroinvasion of intranasal VEEV involves early infection and anterograde transport from immature OSNs. **A-B)** Immunostaining of murine ONE following intranasal VEEV-ZPC-GFP infection (50 pfu, 1 dpi). Solid and hollow arrowheads indicate GFP-labeling of infected GAP43+ immature OSNs (red) and OMP+ mature OSNs (blue), respectively. **(B)** GFP-labeling of OMP+ and GAP43+ OSN axons in the ONE and outer nerve layer of the olfactory bulb following ZPC-GFP infection (50 pfu, 1 dpi). **C)** FlSH staining of VEEV genome (magenta) within the OMP+ (red) and GAP43+ (green) OSN axons in the ONE and outer nerve layer of the olfactory bulb (ZPC-738, 10 pfu, 1 dpi). D) Cartoon depiction of sagittal section nasal cavity and forebrain. ONE (cyan), OB (green), olfactory tract (dark green) are indicated. **E)** Immunostaining of intranasal VEEV-ZPC-GFP infection (50 pfu, 3 dpi) of sagittal sectioned nasal cavity and forebrain, and rostal sequence of coronal slices (Bregma ∼2.0 and -2.0 mm) depicting VEEV-GFP dissemination along lateral olfactory tract and piriform cortex. All images depict representative infections from N=3 mice.

### Higher levels of expression of a VEEV receptor, *Ldlrad3*, underlies enhanced tropism to iOSN

Despite the knowledge that peripheral neurons, including OSNs, are targets for many neurotropic viruses, there are few studies reporting their differential expression of viral entry receptors and innate immune responses [18–20]. To determine whether the SARS-CoV-2 entry receptor angiotensin converting enzyme (ACE2) is an ISG within cells of the ONE, Ziegler et. al. performed single-cell RNA sequencing of murine nasal epithelium derived from mice 12 hours after i.n. administration of saline versus IFN*α* (10^4^ U) [21]. While they found little to no ACE2 expression in iOSN or mOSN (with our without IFN*α* exposure), they provided a large dataset for investigation of the differential expression of other viral entry receptor mRNAs and overall innate immune response networks in iOSN and mOSN [22]. To define differences between transcriptomic signatures of iOSN and mOSN that would underlie the observed enhanced infectivity of VEEV to iOSN, we analyzed differentially expressed genes (DEG) between the two cell types under saline treatment. As expected, the top DEG included genes involved in olfactory sensory perception, cilium development, and ion channel/transport protein expression (Adcy3, Omp, Pde1c, Cngb1, Cnga4 (S2A Fig) [23,24]. Similarly, gene set enrichment analysis (GSEA) using murine gene ontology (GO) pathways identified key differences in pathways associated with neuronal differentiation, axonal growth and synapse formation between iOSN and mOSN (S2B Fig). While mRNAs of genes relating to innate immunity or control of virus infection were not among the top DEG, some genes associated with GO pathways, including Innate Immune Response, Response to Virus, Response to Type 1 Interferon, Response to Interferon Alpha, and Response to Interferon Beta, were differentially expressed between iOSN and mOSN (Fig 2A). For example, mRNA levels of Interferon Induced Protein With Tetratricopeptide Repeats 1 (*Ifit1*) was significantly higher in mOSN (Fig 2B). However, none of these pathways were more significantly enriched by GSEA in either OSN population (Table 1), consistent with lack of differences in mRNA expression levels of IFNαβ receptor (*Ifnar1*) (Fig 2B, S2B Fig). Together, broad differences in innate immunity do not explain the observed early VEEV tropism and infection of iOSN compared with mOSN. However, mRNA levels of a VEEV receptor, *Ldlrad3* [4], are significantly higher in iOSN versus mOSN (Fig 2B). While detection of Ldlrad3 mRNA via fluorescent in situ hybridization (FISH) was observed in both iOSN and mOSN (Fig 2C), it is likely that overall difference in levels of expression of *Ldlrad3* underlie earlier infection of iOSN.

**Fig 2.**
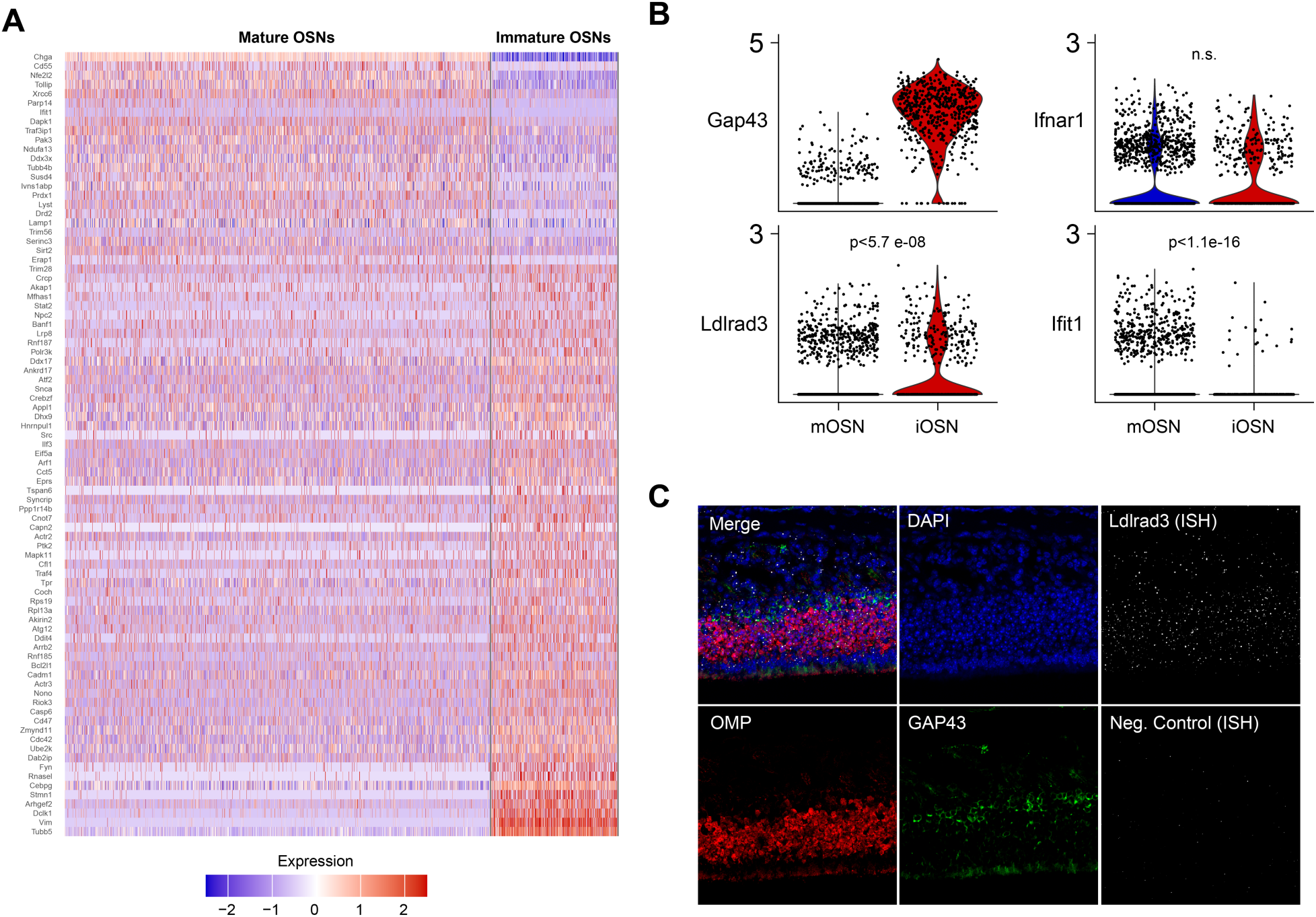
Differential expression of innate immune genes and LDLRAD3 in OSN **A)** Reanalysis of scRNA-seq dataset (Ziegler et al., 2020) which captured a large population of both immature and mature murine OSN. Heatmap of DEGs belonging to GO terms (Innate Immune Response, Response to Virus, Response to Type-1 Interferon, Response to Interferon Alpha, and Response to Interferon Beta (N=1816 mOSN, 539 iOSN from two mice). **B)** Violin plot of expression of candidate genes relevant to VEEV infection and interferon signaling in mature and immature OSN. C) FISH of Ldlrad3 (gray) expression in OMP+ (red) and GAP43+ (green) OSNs within the ONE. Images depict representative staining from N=3 mice. P values are indicated.

**Table 1.** GSEA analysis of candidate innate immune GO pathways in mOSN and iOSN.

| **GO Pathway** | **ID** | **Adj. P Val** | **NES** |
| --- | --- | --- | --- |
| Innate Immune Response | GO:0045087 | 0.68 | 1.081 |
| Response to Virus | GO:0009615 | 0.69 | 1.082 |
| Response to Type 1 Interferon | GO:0034340 | 0.99 | 0.539 |
| Response to Interferon Alpha | GO:0035455 | 0.94 | -0.683 |
| Response to Interferon Beta | GO:0035456 | 0.80 | 0.99 |

### Intranasal IF**N**α administration induces rapid ISG expression in olfactory sensory neurons

Given that OSN exhibit low levels of expression of innate immune molecules at baseline, we analyzed the scRNAseq dataset deposited by Ziegler et. al. for DEG in iOSN and mOSN following intranasal IFNα (10^4^ U, 12 h) treatment [21,22]. Both OSN cell types exhibit similar DEG (Fig 3A-B). As expected DEGs and GSEA indicate strong enrichment of ISGs relating to Type 1 interferon responses following treatment, which was broader for mOSN (S2C-D Fig). Overall, this indicates that these cells, critical to early ONE replication and neuroinvasion into the OB, are responsive to such treatment. To validate ISG expression in the ONE and OB in a separate cohort of mice, we examined candidate ISG expression in total nasal cavity (NC) and OB following similar i.n. administration of recombinant murine IFNα (8×10^4^ U) followed by quantitative (q)PCR. Robust upregulation of ISG, including *Ifit1, Irf7, Ifitm3, Isg20, and cGas,* mRNAs was observed in both the nasal cavity and OB at 24 hours post-treatment with IFNα compared with vehicle-treated animals (Fig 3C). To determine if IFNα if also altered expression of Ldlrad3, we quantified *Ldlrad3* expression as assessed by FISH within the ONE following IFNα treatment at 12 hpi during VEEV infection (Fig 3D). While *Ldlrad3* expression was enhanced by VEEV infection, IFNα treatment did not synergistically impact its level of expression. Overall, these data indicate that IFNα treatment elicits rapid IFN response in both the ONE and OB, suggesting a potential therapeutic approach for limiting infection and neuroinvasion along this route.

**Fig 3.**
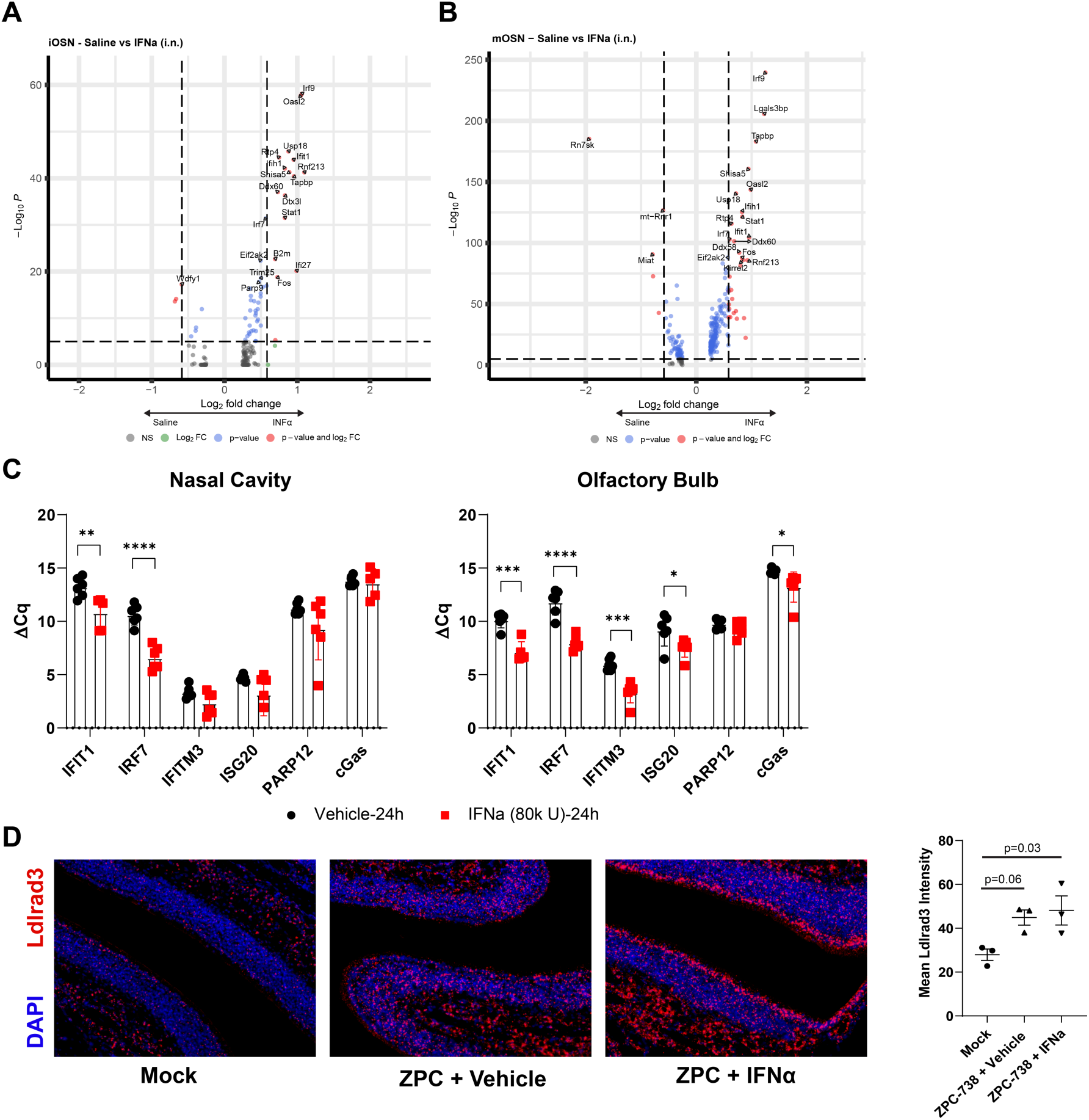
Single-dose intranasal IFNα treatment stimulates ISG expression in OSNs and OB. **A)** Volcano plot of DEGs within immature OSNs following intranasal saline (negative) or IFNα (positive) treatment (1×10^4^ U, 12 hours, N=539 saline, 561 IFNα from two mice). **B)** Volcano plot of DEGs within mature OSNs following intranasal saline (negative) or IFNα (positive) treatment (1×10^4^ U, 12 hours, N= 1816 saline, 2076 IFNα from two mice). **C)** Representative ISG expression within nasal cavity and olfactory bulb homogenates following intranasal IFNα (8×10^4^ U, 24 h, N=5-6) treatment. **D)** FISH of Ldlrad3 expression vehicle or IFNα co-administered alongside intranasal VEEV ZPC (10 pfu, 12 hpi, N=3). Error bars indicate mean ± SEM. ΔCq were compared via Student t-test. Ldlrad3 expression was compared by one-way ANOVA, followed by Tukey multiple comparison. Statistical values are indicated as follows *, *P*<0.05; **, *P*<0.01; ***, *P*<0.001, ****, P<0.0001 unless otherwise stated.

### VEEV-mediated induction of endogenous IFN and ISGs is delayed within infected olfactory routes

While endogenous IFN signaling is critical for controlling VEEV infection in the periphery [7], the extent to which it controls VEEV infection and dissemination along the olfactory route is unknown. Knowledge of the kinetics of this response is also important for determining if exogenous administration of IFNα would be expected to limit VEEV neuroinvasion. To address this, IFN mRNA expression within the nasal cavity and OB was assessed in uninfected animals and at various time points (12, 24, 48 hours) post-infection (hpi). IFNα mRNA is not significantly upregulated in the nasal cavity or OB until 48 hpi (Fig 4A), while IFNβ mRNA induction is observed at 12 and 24 hpi within the nasal cavity, with significant induction at 24 hpi in the OB. Separate analyses of ISG mRNAs linked to inhibition of alphavirus infection showed similarly delayed onset of expression in the nasal cavity and OB (Fig 4C-H). Only IRF7 was upregulated within 24 hpi (Fig 4D-E), with all other candidates not exhibiting expression in both the NC and OB until 48 hpi (Fig 4C-H). These data indicate potential windows of intervention with i.n. administered IFNα after i.n. exposure to VEEV.

**Fig 4.**
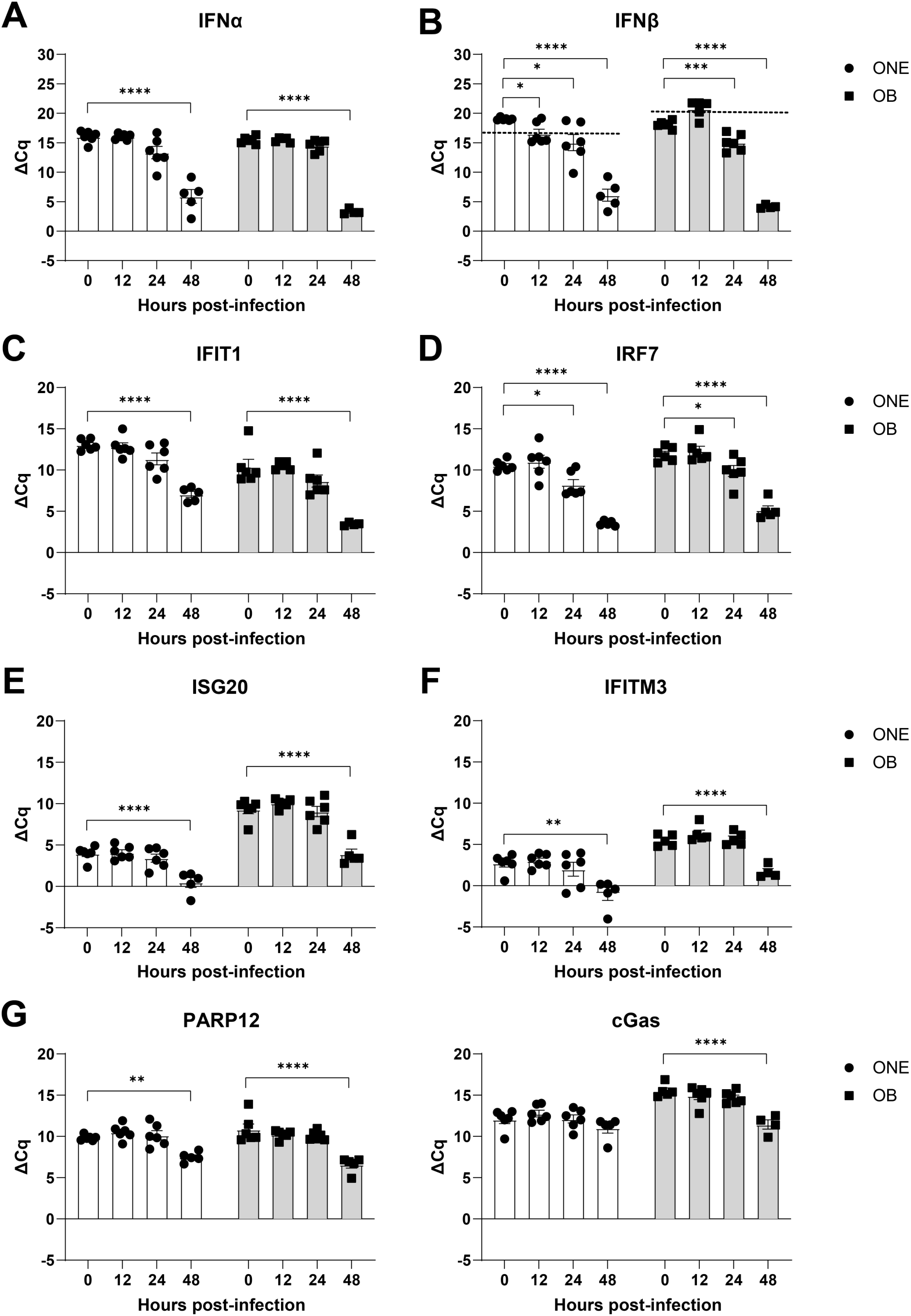
Interferon response in nasal cavity and OB is delayed relative to infection. **A-B)** Induction ONE and olfactory bulb (OB) expression of endogenous type-I interferons, IFNα **(A)** and IFNβ **(B)** within nasal cavity and olfactory bulb homogenates following intranasal inoculation of VEEV-ZPC-738 (10 pfu). **C-H)** Upregulation of candidate interferon stimulated genes (ISGs) associated with restricting VEEV and/or alphavirus replication induced nasal cavity and olfactory bulb homogenates at various time points following intranasal inoculation of VEEV-ZPC-738 (10 pfu). Error bars indicate mean ± SEM, N=5-6. ΔCq were compared via one-way ANOVA, followed by Dunnett’s multiple comparison test. Statistical values are indicated as follows *, *P*<0.05; **, *P*<0.01; ***, *P*<0.001, ****, P<0.0001 unless otherwise stated.

### Intranasal IFNα treatment early after VEEV exposure delays morbidity and promotes survival

To determine if i.n. administration of IFNα during early VEEV infection would improve outcomes following VEEV infection, IFNα treatment (8×10^4^ U) was administered concomitantly or at 1 and 3 hours after i.n. VEEV infection (ZPC-738, 10 pfu) of wild-type mice. Pre-treatment (0 hpi) with IFNα delays morbidity, as assessed via encephalitic scoring and weight loss, compared with similarly infected vehicle-treated mice (Fig 5A). Specifically, weight loss was significantly lower in IFNα-treated mice, and onset lagged approximately 2 days behind that of vehicle-treated VEEV-infected mice (Fig 5A, left). While VEEV encephalitic signs were similar between vehicle- and IFNα-treated animals, scores were significantly lower and delayed by approximately 2 days in IFNα-treated mice (Fig 5A, middle). IFNα-treatment at the time of VEEV infection extended mean survival time (+ 2 dpi MST) but was ultimately insufficient to improve overall mortality (Fig 5A, right). As knowledge of exposure to VEEV may be delayed, we determined whether post-infectious IFNα treatment at 1 or 3 hpi impacts disease and survival after i.n. infection with VEEV. Both treatment paradigms significantly delayed onset of encephalitic sequelae and weight loss compared with vehicle-treated VEEV-infected mice (Fig 5B, left and middle). However, weight was not as well maintained as observed during concomitant IFNα and VEEV i.n. exposure compared with similarly infected vehicle-treated mice, especially when IFNα was administered at 3 hpi. Similar to concomitant treatment, IFNα-treatment extended mean survival time (+2 dpi and +1.6 dpi, respectively), without reducing overall mortality (Fig 5B, right).

**Fig 5.**
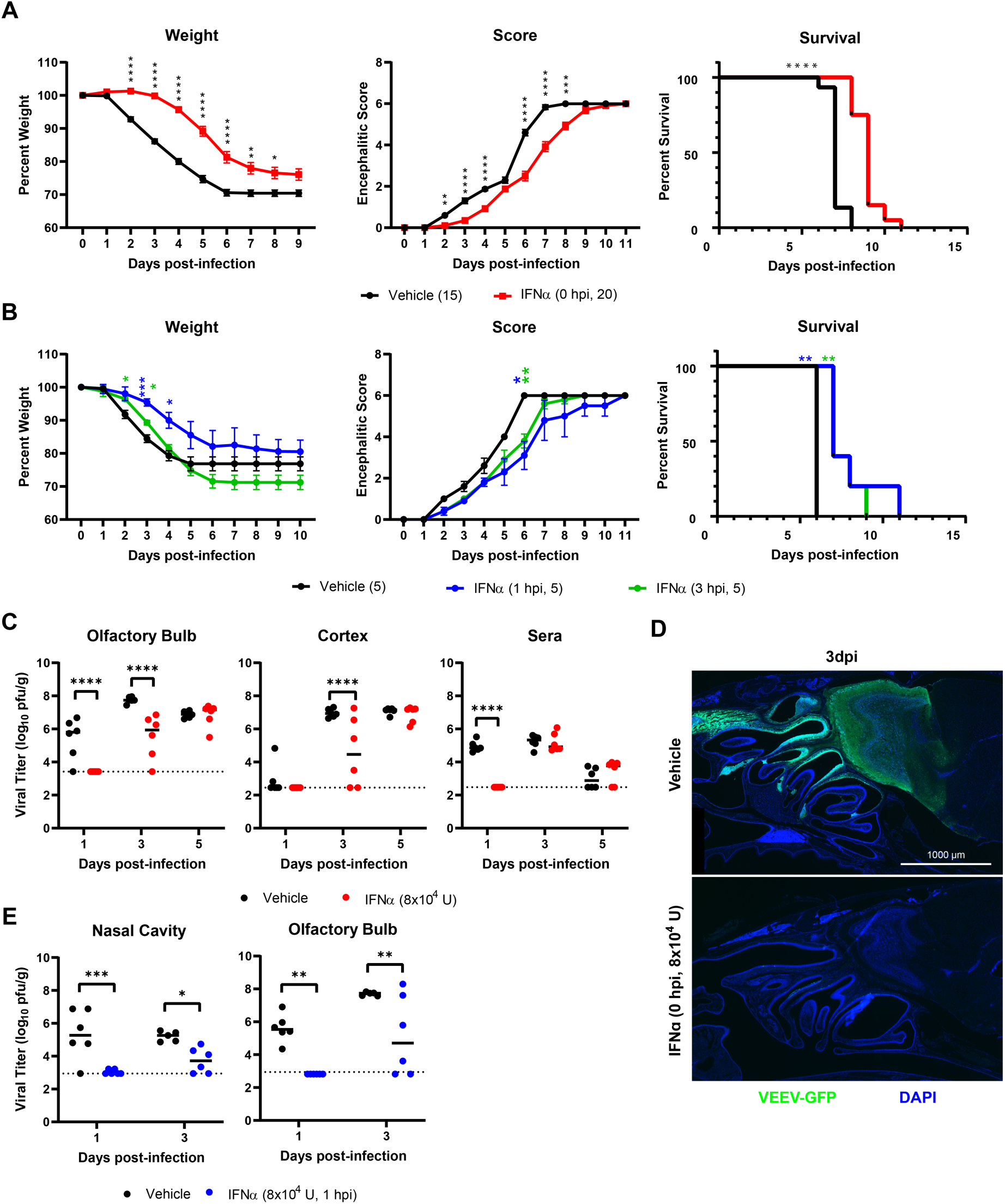
Intranasal IFNα delays morbidity and promotes survival during VEEV infection by suppressing onset of nasal and CNS infection. **A)** Survival curves, weight loss curves, and encephalitis scores of mice co-administered single-dose intranasal IFNα during ZPC-738 (10 pfu) infection (N=15-20 mice from two independent infections). **B)** Survival curves, weight loss curves, and encephalitis scores of mice administered single-dose intranasal IFNα 1 or 3 hour post-infection with VEEV ZPC-738 (10 pfu) (N=5). **C)** Viral titers, as measured by plaque assay, from olfactory bulb, cortex, and serum collected at various DPI from mice co-administered intranasal IFNα during ZPC-738 (10 pfu) infection (N=6). **D)** Immunostaining of intranasal VEEV-ZPC-GFP infection (50 pfu, 3dpi) of sagittal sectioned nasal cavity and forebrain (Representative example for N=3 mice). **E)** Viral titers from nasal cavity and olfactory bulb at various DPI to following intranasal IFNα administered 1 hour following ZPC-738 (10 pfu) infection (N=5—6). Weight loss, encephalitic sequelae scores, and titers were compared via two-way repeated measure ANOVA, followed by Bonferroni’s post hoc test. Survival curves were analyzed by Mantel-Cox test. Statistical values are indicated as follows *, *P*<0.05; **, *P*<0.01; ***, *P*<0.001, ****, P<0.0001 unless otherwise stated.

To determine if i.n. administration of IFNα limited VEEV replication within the ONE and/or CNS dissemination, viral titers were assessed in IFNα-treated animals. For mice administered IFNα at the time of VEEV infection (0 hpi), viral titers were assessed at 1, 3, and 5 dpi via standard plaque assays. In contrast with vehicle-treated animals, viral titers were undetectable in the brain (OB, CTX) or sera in IFNα-treated mice at 1 dpi (Fig 5C). By 3 dpi, VEEV was detectable in the majority of CNS tissues derived from vehicle- and IFNα-treated mice (VEEV OB: 5/6; CTX: 4/6). However, overall VEEV titers in OBs and cortices derived from IFNα-treated mice were significantly reduced at this time-point compared with similarly infected vehicle-treated animals (Fig 5C). IFNα treatment, however, failed to control viremia beyond 1 dpi, as viral titers were equivalent in both treatment groups of VEEV-infected mice by 3 dpi. Similarly, VEEV viral titers in IFNα-treated mice reached equivalency to vehicle-treated mice by 5 dpi in all CNS regions (Fig 5C). Direct observation of i.n. ZPC-738-GFP infection (50 pfu) in fixed whole skull mounts revealed strong suppression of infection in the ONE and OB at 3 dpi in IFNα-treated compared to vehicle-treated mice, which displayed robust ZPC-738-GFP throughout the ONE, OB, and olfactory tract (Fig 5D). Finally, similar to concurrent (0 hpi) administration, i.n. IFNα treatment after infection (1 hpi) suppressed early infection, as VEEV viral titers at 1 dpi within the nasal cavity (NC) and OB were undetectable compared with vehicle-treated mice (Fig 5E). By 3 dpi, infection in the nasal cavity and OB remained significantly reduced in VEEV-infected mice treated with IFNα compared with vehicle-treated animals but was detectable in a majority of animals (VEEV+ NC: 4/6, VEEV+ OB: 4/6).

## DISCUSSION

In this study, we tracked the spread of VEEV infection from the ONE to the ONL of the OB, examining viral targets, innate immune responses, and the efficacy of post-exposure treatment with i.n. IFN. We found that GAP43+ iOSN were the first cells infected, followed by OMP+ mOSN, which both transport VEEV anterograde into the OB. scRNAreq analysis of OSN identified no broad innate immune deficits associated with iOSN to explain their enhanced infectivity compared to mOSN. However, specific changes in key ISGs, including significantly decreased levels of IFIT1 and/or increased expression of the VEEV receptor LDLRAD3 may underlie VEEV tropism to iOSN. The kinetics of ISG expression after i.n. VEEV revealed a significant delay, with robust upregulation occurring after 24 hours. To determine if ISG levels could be rescued by exogenously administered IFN we utilized a model of i.n. IFNα treatment at the time of or post-exposure to VEEV infection. We found that IFNα treatment triggers early ISG expression in OSN, the nasal cavity and OB, even when administered as a single post-exposure dose. Consistent with this, IFNα treatment delayed onset of VEEV infection in the nasal cavity and OB, reduced encephalitic sequelae and extended survival. These data demonstrate that exogenous IFNα may be a potential post-exposure intervention for VEEV infection, allowing infected individuals time to obtain additional support or other treatments.

In concordance with our findings, previous studies have demonstrated VEEV infection of OSN, however these studies did not distinguish tropism between iOSN versus mOSN [25,26]. Axonal transport of VEEV has also been previously described, with detection of VEEV antigen and virions in olfactory nerve fibers crossing the cribriform plate [26]. Depending on their stage in maturation, iOSNs fully project to OB by ∼7 days of differentiation, forming functional synapses in OB glomeruli that participate in limited olfaction [27–29]. As these neurons continue to express markers of immaturity, iOSNs represent not only an early site of VEEV infection within the ONE but also a route of anterograde transport to the OB. As VEEV infection propagates within the ONE, infected OMP+ mOSNs likely also contribute to additional VEEV anterograde transport of VEEV to OB but may be less critical to initial neuroinvasion along the olfactory tract.

Examination of genetic signatures of iOSN and mOSN at baseline and after IFN exposure was performed via interrogation of a previously deposited scRNAseq dataset [21,22]. Type-1 IFN induces expression of ISGs, with only a few mediating the anti-viral activity for a specific pathogen. We found no broad deficits in innate immunity or antiviral gene expression at baseline to explain enhanced infectivity of iOSN over mOSN, with the exception of *Ifit1,* which was more highly expressed in mOSN. IFIT1 has been shown to limit VEEV replication by restricting translation of VEEV in strains that contain a G3A mutation, such as the TC-83 vaccine strain [30]. However, IFIT1 may be involved in other mechanisms that restrict VEEV replication since, within the same study, Ifit1-/-mice exhibited shorter MST for both WT ZPC-738 and TC-83 (A3G) mutants. Additionally, IFIT1 positively enhances ISG expression independently of viral RNA binding downstream of TLR4 activation in macrophages [31]. It is also possible that OSN differentiation induces other protective effects. Neuronal differentiation was observed to restrict VEEV infection *in vitro* using the AP7 olfactory-derived neuronal cell line [32]. This effect was cell intrinsic for differentiated cells and correlated with enhanced expression of interferon response factor (IRF)-3 and -7. Thus, additional screening of identified genes might be warranted. Most notably, mRNA expression of the VEEV receptor *Ldlrad3* was enhanced in iOSNs compared to mOSN. The endogenous ligand for LDLRAD3 is unknown, and is proposed to be distinct from other LDL receptor family members [33]. The role of LDLRAD3 in the maturation of iOSN is unknown, however LDLRAD3 modulates amyloid percursor protein in neurons and promotes activity of E3 ubiquitin ligases, both of which impact neurogenesis [33–36].

Intranasal Type 1 IFN therapy has been explored for various respiratory viruses, including endemic viruses (rhinovirus and influenza) and recently SARS-CoV2 to modulate the severity of disease [37]. However, our student presents the first demonstration of IFN administration to treat a potential aerosolized biological weapon. Encephalitic alphaviruses have evolved immune evasion mechanisms that inhibit host IFN responses and allow virus replication in infected cells. As mentioned above, IFN signaling is suppressed by global shut-off of host transcription and translation and inhibition of STAT-1 signaling by capsid and capsid-independent mechanisms [8,9,9,12]. However, our study demonstrates that the nasal cavity, including OSNs, responds rapidly to intranasal administration of IFN with detectable changes of ISG expression within 12 hours, overcoming IFN restriction by VEEV. The antiviral state initiated following early IFN treatment after VEEV infection leads to suppression VEEV replication in the nasal cavity, preventing early expansion of VEEV infection and escape of VEEV into the blood. Previous studies have shown similar transient protection following intranasal VEEV infection, although these studies utilized prophylactic, multiday treatment and pegylation of IFNα (i.p.) [38]. However, IFN treatment is not able to control VEEV CNS infection indefinitely. It remains unclear from which reservoir VEEV re-emerges after the effects of exogenous IFNα waned. It is possible that additional peripheral sites did not receive sufficient exogenous IFNα to fully prevent VEEV infection, allowing for infection of the ONE and OB via hematogenous routes. Intranasal delivery has been shown to enhance IFN delivery to rodent and non-human primate brain, especially the OB [14,15]. Consistent with this model, we observed ISG expression in the OB of treated mice and sustained suppression of OB viral titers in the presence of normal viremia at 3 dpi. This may indicate additional protection of CNS infection downstream of ONE infection. We’ve reported previously that OB is also an early site of VEEV CNS infection following subcutaneous infection [3]. Models utilizing subcutaneous or intravenous inoculation could potentially elucidate whether protection of OB is due to local IFN signaling or predominately secondary to delayed replication in the ONE.

Overall, intranasal IFN treatment delays onset of morbidity and extension of survival in a highly lethal animal model of VEEV infection. ZPC-738 is an enzootic strain that is completely lethal in mice. Such a disease course does not reflect the lethality associated with VEEV infection in humans. Approximately, >1% of patients with VEEV succumb to the infection [1,39]. Therefore, the IFN treatments strategies explored herein may yet be more effective in protecting against lethal VEEV encephalitis in patients. Certainly, early intervention will likely be the most effective, but additional studies evaluating sustained and late IFNα treatment are warranted. Future studies may continue to leverage murine models to explore IFN modification and delivery strategies. Pegylation of type-1 interferon sustains bioavailability, and improved outcomes against VEEV when administered i.p. [38]. Other modifications focusing on enhancing retention in the nasal cavity with modification or in situ mucoadhesive gel solution [40,41]. However, modifications would need to be evaluated for ease of delivery to the ONE, the effect on IFN delivery to CNS, and how VEEV neuroinvasion would be impacted, especially in those strategies that may disrupt the nose-to-brain barriers [42].

## MATERIAL AND METHODS

### Animals

C57BL/6J mice were purchased from Jackson Laboratories (Bar Harbor, ME). Animals were housed under pathogen-free conditions in Washington University School of Medicine animal facilities. All experiments were performed in compliance with Washington University animal studies guidelines.

### Mouse model of VEEV encephalitis

8-10 week old male mice were inoculated intranasally (10 µL per nostril) with VEEV strain ZPC-738 or ZPC-738-eGFP (10 PFU or 50 PFU, respectively) under anesthesia. ZPC-738-eGFP was a generous gift of William Klimstra (Pittsburg, PA). ZPC-738-GFP was generated by subgenomic insertion of GFP as a cleavable element between the capsid and PE2 structural proteins, described previously (Sun et al., 2014). Mice were monitored daily for weight loss and scored daily for encephalitic sequelae. Moribund mice were sacrificed by CO_2_ asphyxiation and recorded as dead the following day. Encephalitic score represents a progressive range of behaviors: 1) hunched, ruffled fur, 2) altered gait, slow movement, 3) not moving but responsive, 4) not moving, poorly responsive but upright, 5) moribund, 6) dead.

### Perfusion-fixation and immunohistochemistry

At various times post-infection, mice were anesthetized followed by extensive cardiac perfusion with PBS and perfusion fixation with 4% paraformaldehyde (PFA) in PBS. Tissue was immersion-fixed for an additional 24 hours in 4% PFA. For slice preparations of mouse nasal cavities, skulls were decalcified by multiple exchanges 0.5 M EDTA (pH 7.4) in PBS over 7 days followed by PBS and cryoprotection (two-exchanges of 30% sucrose for at least 48 hours) and embedding in OCT (Fisher). 10 μm-thick fixed-frozen sagittal sections were hydrated with PBS and blocked for 1 h in blocking solution, 5% normal donkey serum (Santa Cruz Biotechnology) with 0.1% Triton X-100 (Sigma-Aldrich). After block, slides were exposed to primary antibody at 4°C overnight, washed with PBS and incubated with Alexa Fluor donkey secondary antibodies (Invitrogen) for 1 h at room temperature. Antibodies used: chicken anti-GFP (Abcam, 13970), goat anti-OMP (Wako Chemicals, 544-10001), rabbit anti-GAP43 (Novus Biologicals, NB300-143). Images were acquired using a Zeiss LSM 880 confocal laser scanning microscope and processed using Zen3.3 (Zeiss) and Image J. Quantification of immunofluorescence was performed using ImageJ.

### In situ hybridization

In situ hybridization staining of decalcified sagittal skull section (described above) were performed using Advanced Cell Diagnostics (ACD) RNAscope system and probes. After rehydration of slides in PBS, slides were baked (30 min at 60°C) and post-fixed in 4% PFA. Slides were dehydrated in progressive ethanol washes (50%, 70%, 100%, 100%, 5 min), air dried, treated with hydrogen peroxide (10 min). For in situ hybridization alone, Advanced Cell Diagnostics RNAscope 2.5 HD Detection Reagent – RED (322360) using standard manufactures protocol, RNAscope Target Retrieval Reagent (95-98°C, 10 min), RNAscope Protease Plus (30 min), and standard hybridization with the Ldlrad3 probe (ACD,), signgal amplification, and counter-staining with DAPI. For combined RNA-protein co-imaging, RNAscope Multiplex Fluorescent v2 Assay (323100) along with RNA-Protein Co-detection Ancillary Kit (323180) was used utilizing the Integrated Co-Detection Workflow (ICW). Following baking, post-fixation, dehydration, and hydrogen peroxide treatment, slides were immersed in Co-Detection Target Retrieval (95-98°C, 5 min). Tissue was blocked and incubated overnight with GAP43 and OMP primary antibodies (see above) using Co-Detection Antibody Diluent.

Samples were post-primary fixed using 10% neutral buffered formalin (30 min, RT) prior to RNAscope Protease Plus treatment, hybridization with the V-VEEV-ZPC-738 (ACD, 876381), Mm-Ldlrad3 (ACD, 872101), or dapB negative control probes (ACD, 310043), signal amplification with Opal 650 Dye (Akoya Biosciences, OP-001005) in RNAscope Multiplex TSA Buffer. Tissues were labeled with Alexa-conjugated secondary antibodies (see above) in Co-Detection Antibody Diluent, counter-stained with DAPI, and mounted in Prolong Gold (Invitrogen #P36930). Tissue were imaged as described above.

### Interferon Treatment of Mouse Nasal Mucosa

scRNAseq dataset of intranasal IFNα mice was originally generated as described previously. Briefly, 8-10 week old C57BL6/J mice received either 200ng of IFNα (Biolegend 752802, ∼ 1×104 U) or saline intranasally (N = 2), Respiratory and olfactory mucosa were isolated 12 h later. Single cells suspension were generated in media containing Liberase (Roche) and DNase I (Roche) and loaded on duplicate Seq-Well S3 arrays for sequencing using Illumina NextSeq. Raw expression counts for cells previously defined within Immature Olfactory Sensory Neurons and Olfactory Sensory Neurons (saline and IFNα treated) clusters were downloaded from published dataset. https://singlecell.broadinstitute.org/single_cell/study/SCP832?scpbr=the-alexandria-project#study-summary. Data was normalized and scaled using the Seurat R package (https://satijalab.org/seurat/). Differential expression tests between mature and immature OSNs within saline-treated group or between saline-treated or IFNα-treated OSNs were performed using Seurat FindAllMarkers function with default settings and Wilcoxon rank sum test (p value threshold = 0.05). GSEA analysis was performed using fgsea function from (fgsea, using the murine Gene Ontology gene sets (MSigDB). Genes were ordered by the Log2 fold change using Seurat FindMarkers function. Violin plots and heatmaps were generated using Seurat R package. Volcano plots were generated using the EnhancedVolcano package.

### Administration of IFNα

Recombinant mouse IFNα (PBL Assay Science, 12100-1) was administered intranasally. Control mice were similarly administered vehicle solution of 0.1% bovine serum album (BSA) in PBS. Doses (8×10^4^ U, 10 µL/nostril) administered at time of infection were suspended in the inoculum under brief isoflurane anesthesia. Subsequent doses (8×10^4^ U, 5 µL/nostril) were administered at 1-3 hours post-infection (hpi), as indicated, under brief isoflurane anesthesia.

### RNA isolation and quantitative RT-PCR

Cortices were isolated from cardiac-perfused mice at 12, 24, and 48 hours after ZPC-738 (10 PFU i.n.). RNA was isolated from tissues using RNeasy kit (Qiagen) according to manufacturer’s instructions, and quantified using a NanoDrop (Thermo Scientific). Following DNAse I treatment (Invitrogen) of RNA samples (1 µg) was reverse transcribed using Taqman Reverse Transcriptase kit (Applied Biosystems). qRT-PCR was performed using Power SYBR Green (Applied Biosystems) on a CFX3384 PCR Detection System (Bio-Rad) using manufacturer’s recommended cycle parameters. Values are reported as the Cq values for target genes normalized to Cq values of GAPDH (Ct_gene_/Ct_GAPDH_). Primers (5’-3’) used are reported in Table S1.

### Virologic analysis

At various post-infection intervals, CNS tissue was collected from ZPC-738 infected after extensive cardiac perfusion with PBS. Viral titers were determined using standard plaque assay techniques by serial dilution of tissue homogenates over BHK cells, as described previously (Brien et al., 2013).

### Statistical analyses

Reported values are mean values ± standard error of the mean (SEM). Statistical analysis was performed using GraphPad Prism 7 software. Survival curves were analyzed by Mantel-Cox test. Cytokine and ISG expression in mice were analyzed via one-way analysis of variance (ANOVA), Bonferroni’s post hoc test was subsequently used for comparison of individual means. Weight loss and encephalitic sequelae scores were compared via two-way repeated measure ANOVA, followed by Bonferroni’s post hoc test. P values *P* < 0.05 were considered significant. Statistical values are indicated as follows *, *P*<0.05; **, *P*<0.01; ***, *P*<0.001, ****, P<0.0001 unless otherwise stated.

## Acknowledgements

We would like to thank Drs. Michael Diamond and Natasha Kafai (Dept. of Medicine, WUSM) for guidance and shared reagents for Ldlrad3 detection, Qingping Wu and Katie Madden (Dept. of Medicine, WUSM) for assistance in early experiments, Wandy Beatty of the Molecular Microbiology Imaging Facility (WUSM) for advice on confocal imaging, and Dr. Mark Miller for input on the original concept.

Funding for this study was provided by R01AI175524 and R35NS122310 (to RSK).

